# Gamma-rhythmic Gain Modulation

**DOI:** 10.1101/060582

**Authors:** Jianguang Ni, Thomas Wunderle, Christopher M. Lewis, Robert Desimone, Ilka Diester, Pascal Fries

## Abstract

Cognition requires the dynamic modulation of effective connectivity, i.e. the modulation of the postsynaptic neuronal response to a given input. If postsynaptic neurons are rhythmically active, this might entail rhythmic gain modulation, such that inputs synchronized to phases of high gain benefit from enhanced effective connectivity. We show that visually induced gamma-band activity in awake macaque area V4 rhythmically modulates responses to unpredictable stimulus events. This modulation exceeded a simple additive superposition of a constant response onto ongoing gamma-rhythmic firing, demonstrating the modulation of multiplicative gain. Gamma phases leading to strongest neuronal responses also led to shortest behavioral reaction times, suggesting functional relevance of the effect. Furthermore, we find that constant optogenetic stimulation of anesthetized cat area 21a produces gamma-band activity entailing a similar gain modulation. As the gamma rhythm in area 21a did not spread backwards to area 17, this suggests that postsynaptic gamma is sufficient for gain modulation.

## Introduction

The flexible modulation of effective connectivity is central to many cognitive functions. Selective attention is a prime example, in which the responses to an attended stimulus are routed forward with enhanced effective connectivity (Reynolds et al., 1999). Enhanced effective connectivity corresponds to an enhanced gain, i.e. a stronger response to a constant stimulus. Two intriguing mechanisms for gain modulation might be provided by neuronal gamma-band synchronization. On the one hand, gamma-band synchronization among presynaptic neurons makes synaptic inputs arrive coincidently at postsynaptic neurons, which increases their postsynaptic impact (Azouz and Gray, 2003; Salinas and Sejnowski, 2001). On the other hand, gamma-band synchronization among postsynaptic neurons entails a characteristic sequence of network excitation followed by inhibition (Atallah and Scanziani, 2009; Buzsáki and Wang, 2012; Salkoff et al., 2015; Vinck et al., 2013), which likely modulates the response to synaptic input. Input that is consistently synchronized to gamma phases with high excitability might benefit from enhanced gain and thereby enhanced effective connectivity, a proposal referred to as the “Communication-through-Coherence” (or CTC) hypothesis (Fries, 2005, 2015).

Gain increases for coincident synaptic inputs have been suggested by mathematical models (Salinas and Sejnowski, 2001). In-vivo intracellular recordings from neurons in the visual cortex of anesthetized cats have demonstrated an adaptive coincidence detection mechanism (Azouz and Gray, 2003). Simultaneous recordings in anesthetized macaque V1 and V2 show that V2 spikes are preceded by coincident V1 spikes (Zandvakili and Kohn, 2015). V1 spike coincidence is provided by gamma-band synchronization, and indeed, V1 spikes occurring at the V1 gamma phase of strongest spiking are most often followed by V2 spikes (Jia et al., 2013). This mechanism likely enhances the impact of attended stimuli. V4 neurons driven by attended stimuli show enhanced gamma-band synchronization (Fries et al., 2001), whose strength predicts the attentional reaction-time benefit on a given trial (Womelsdorf et al., 2006).

Mathematical models have also supported the idea that gamma-band synchronization among postsynaptic neurons rhythmically modulates their gain, such that input consistently arriving at high-gain phases benefits from enhanced effective connectivity (Börgers and Kopell, 2008). Simultaneous recordings at multiple sites within or across visual areas of awake cats and macaques demonstrates that effective connectivity, indexed by power covariation, is systematically modulated by the phase relation between respective local gamma rhythms (Womelsdorf et al., 2007). Similarly, in anesthetized macaque V1, directed influences between recording sites are modulated by the respective gamma phase relation (Besserve et al., 2015).

The gain enhancement through synchronization between pre- and post-synaptic neurons might subserve the selective routing of attended stimuli. Neurons in macaque V4 are selectively entrained by the gamma rhythm of V1 inputs representing the attended stimulus (Bosman et al., 2012; Grothe et al., 2012). For this selective entrainment to cause enhanced effective connectivity, the V4 gamma has to modulate gain rhythmically, as a function of gamma phase. This has been a core requirement of the CTC hypothesis (Fries, 2015), but experimental evidence has so far been lacking. The definitive test for gain modulation by postsynaptic gamma phase uses externally timed test inputs placed at different gamma phases. Such test inputs have been used in two seminal studies that probed consequences of optogenetic pulse trains driving fast-spiking interneurons in mouse somatosensory cortex. When the local neuronal population was entrained by a 40 Hz pulse train, its response to a stimulation of a vibrissa was modulated by the 40 Hz phase at which the stimulus was delivered (Cardin et al., 2009). This rhythmic modulation of neuronal responses also impacts behavior, as shown in a subsequent study that used the same approach in barrel cortex of mice performing a tactile detection task. Detection of low-salience stimuli was improved when input to the optogenetically entrained cortex coincided with high excitability phases (Siegle et al., 2014).

If visually induced gamma in V4 exerted similar gain modulation effects on externally timed test inputs, the abovementioned selective inter-areal gamma-band synchronization for attended stimuli might indeed implement enhanced effective connectivity during visual attention. Here, we present evidence from two experiments, one combining electrophysiology with behavioral analysis in awake macaque visual cortex and a second combining electrophysiology with optogenetic stimulation in anesthetized cat visual cortex. In macaques, we recorded multi-unit activity (MUA) and local field potentials (LFP) in area V4, while a visual stimulus induced a sustained gamma rhythm. At a random time, we changed stimulus color, which gave a change-related firing rate response. We found that the magnitude of this response depended on the V4 gamma phase at which the change-related input to V4 occurred. The gamma-phase dependent response modulation went substantially beyond an additive superposition of a constant response on ongoing gamma-modulated firing, demonstrating the modulation of multiplicative gain. The same gamma phase that led to maximal firing rate responses also led to shortest behavioral reaction times, suggesting that the effect has direct functional relevance. As visual stimulation induces partly synchronized gamma-band activity across ventral visual areas (Bastos et al., 2015a; Bosman et al., 2012; Grothe et al., 2012; Jia et al., 2013; Roberts et al., 2013), this effect could emerge at any stage. To test whether gamma in a higher visual area was sufficient to generate the effect, we used optogenetics in anesthetized cats. Constant optogenetic stimulation of area 21a, the cat homologue of macaque V4 (Payne, 1993), induced sustained gamma-band activity in area 21a, that did not spread to area 17, the homologue of V1. When a visual stimulus was presented at random gamma phases, the phase at which the change-related input to area 21a occurred, modulated the stimulus response, suggesting that an isolated post-synaptic gamma rhythm is sufficient to generate a multiplicative gain modulation.

## Results

### Visually induced gamma rhythm modulates gain

MUA and LFP were recorded from two to four electrodes simultaneously in area V4 of two macaques performing an attention task. Visual stimulation with a patch of grating in the RFs of the recorded neurons induced clear enhancements of V4 MUA rate (Figure 1A, B) and LFP gamma power (Figure 1C, D). At a random time between 0.5 and 5 s after stimulus onset, the grating in the RFs changed from black/white to black/yellow. We analyzed the trials in which the stimulus in the RFs was behaviorally relevant and in which the monkey responded correctly, i.e. in which the stimulus change in the RFs triggered a behavioral response. We found that the stimulus change induced a substantial firing rate response (Figure 1B). Preceding the stimulus change, the visually induced gamma-band response was sustained (Figure 1D). The stimulus onset evoked a transient, time-locked LFP component visible in the time-domain LFP average, the event-related potential (ERP) (Figure 1E). Preceding the stimulus change, the ERP was flat (Figure 1F), because changes occurred at random times between 0.5 and 5 s after stimulus onset.

**Figure 1.**
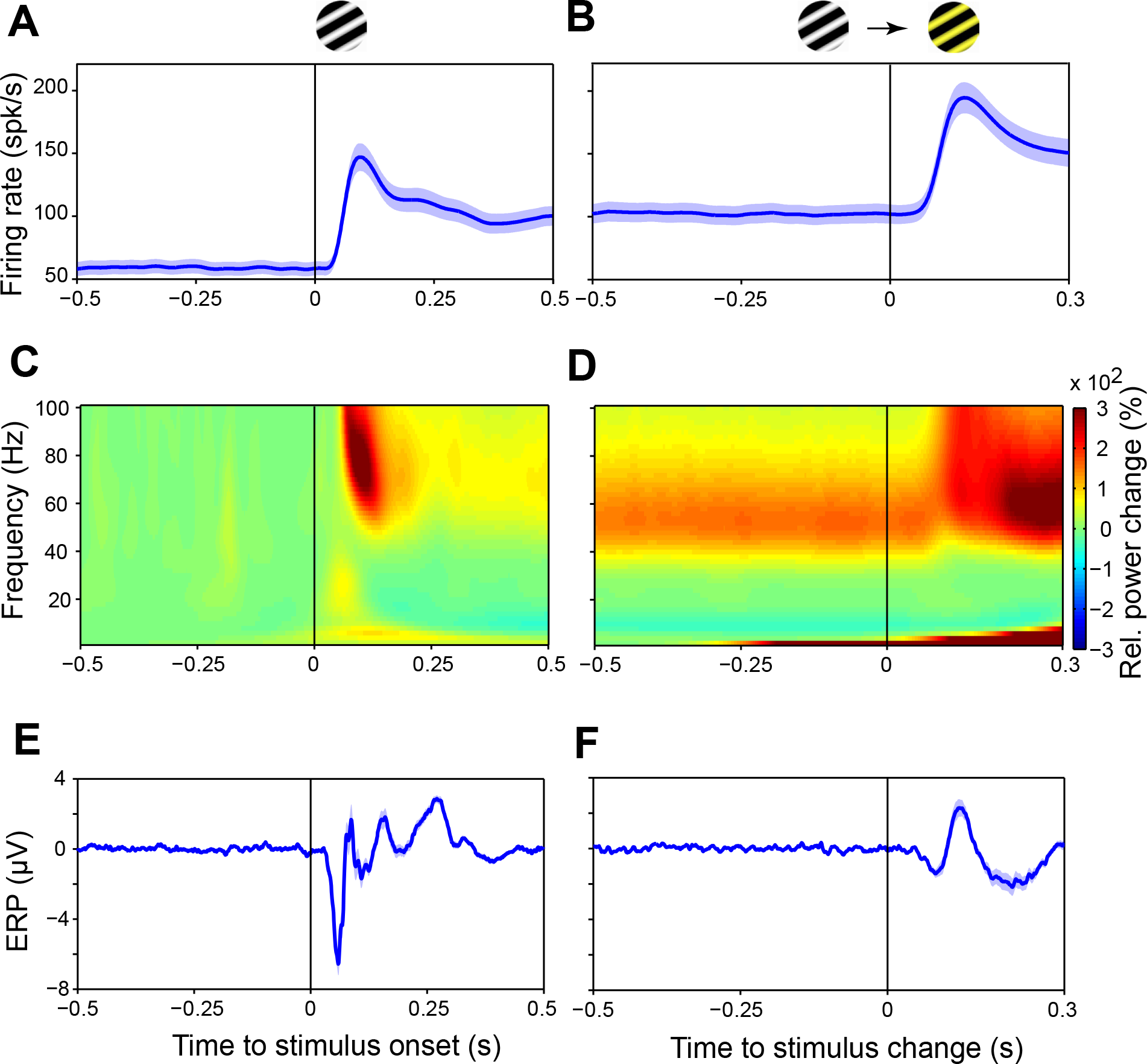
Peri-event Modulation of MUA Firing Rates, LFP Power and Eventrelated Potentials in awake macaque V4. (A and B) MUA firing rate, smoothed with a Gaussian kernel (SD = 12.5 ms, truncated at ±2 SD). (C and D) Percentage LFP power change relative to the pre-stimulus baseline period from 0.5 to 0.25 s before stimulus onset. (E and F) Event-related potentials (ERPs), i.e. time-domain averages of the LFP across trials. (A, C, E) Temporal modulation around stimulus onset. (B, D, F) Temporal modulation around stimulus color change. (A-F) All plots show grand averages over all sites in both monkeys. (A, B, E, F) Shaded regions around the lines indicate ±1 SEM across recording sites.

This allowed us to analyze the ongoing LFP phase and test whether it predicts the MUA response to the stimulus change. Figure 2 illustrates this for an example recording site. We first estimated the time of arrival of the change-induced synaptic inputs, by calculating the inter-trial coherence (ITC) of the LFP (Figure 2A). ITC quantifies the phase locking, across trials, to the stimulus change time. For the example site, the ITC significantly exceeded a shuffle control at 42 ms. This ITC increase constituted the earliest sign of a neuronal response to the stimulus change and was therefore defined as “input time”. The median (±SEM) input times for the two monkeys were 41±2 ms (Monkey P) and 43±4 ms (Monkey R). We defined the LFP phase estimated for the last (1 ms) sample before the input time as the preinput phase (see Experimental Procedures for details).

**Figure 2.**
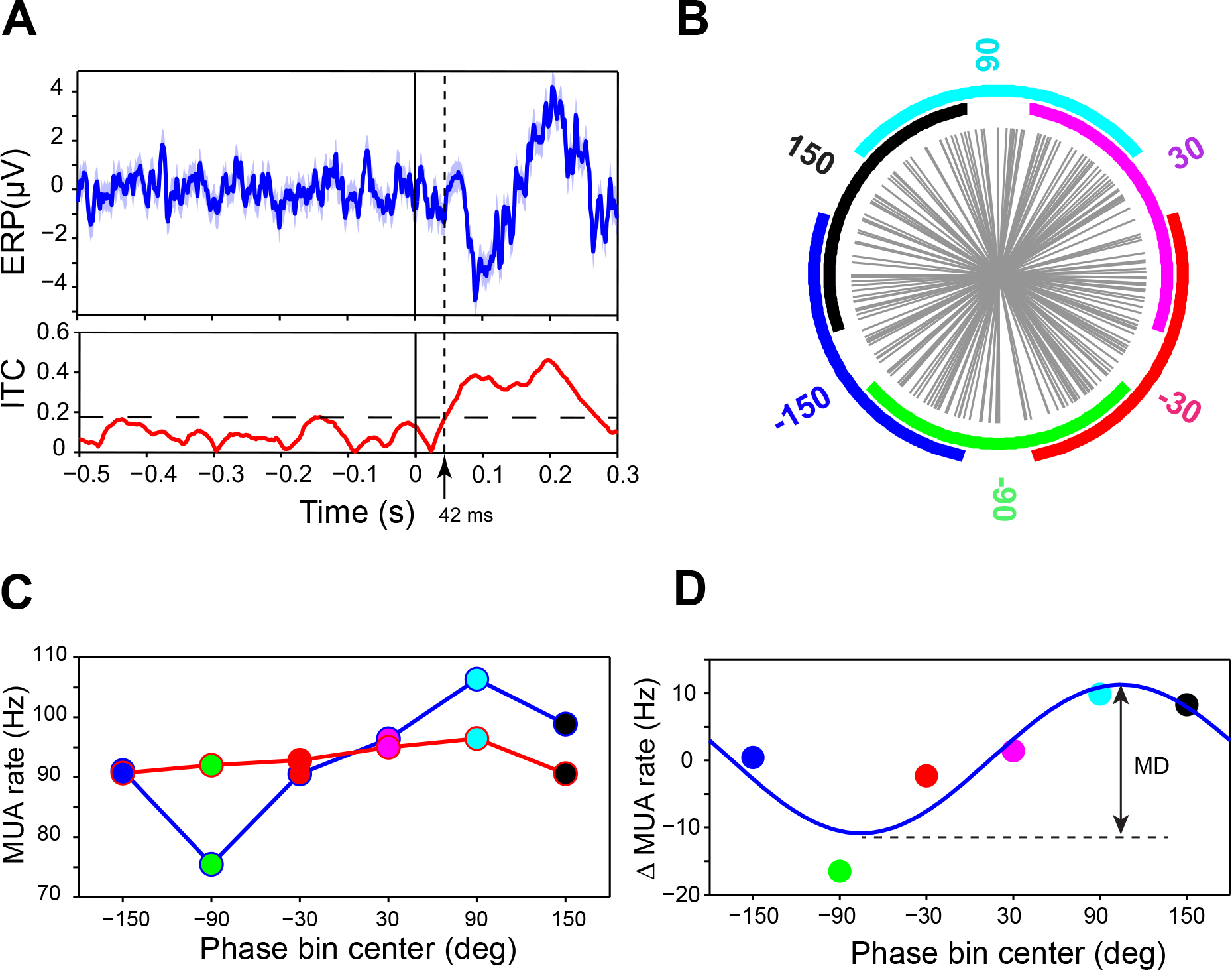
Example Analysis of Response Modulation by Pre-input Phase. (A) The top panel shows the ERP that was evoked in an example recording site by stimulus changes. The bottom panel shows the corresponding inter-trial coherence (ITC) together with the significance threshold, indicating the first change-evoked response at 42 ms after stimulus change. (B) Each gray spoke represents the pre-input LFP phase for the gamma band (50 Hz) in one trial (see Experimental Procedures for details of phase estimation). Trials were grouped into six phase bins. For each phase bin, the 75 trials with phases closest to the phase-bin center were chosen for further processing. (C) Blue line: MUA response as a function of pre-input gamma phase, both parameters averaged per phase bin. Red line: Same as blue line, but showing the additive MUA response component. (D) Colored dots: Multiplicative MUA response component, obtained by subtracting the additive MUA response component from the (total) MUA response. The smooth blue curve represents a cosine fit. The cosine modulation depth (MD) is quantified as indicated.

Pre-input phase distributions were expectedly random, and we binned phases into six bins as indicated by the colored sectors in Figure 2B for the gamma-phase distribution. MUA responses to the stimulus change were quantified for the time, when the trial-averaged MUA peaked, which we call the “peak time”. For the example MUA, the peak time was at 134 ms after stimulus change. MUA responses depended systematically on pre-input gamma phase (blue line in Figure 2C). The dependence of the MUA response on pre-input gamma phase might be due to a simple additive superposition of a constant MUA response onto ongoing gamma-modulated MUA firing; note that MUA is typically synchronized to the visually induced LFP gamma rhythm (Fries et al., 2008). If the MUA response peak would coincide with a peak of gamma-modulated MUA firing, the response would be enhanced, and vice versa. The size of such an additive superposition effect can be estimated by mathematically adding the average MUA response to the pre-input MUA record after phase binning (van Elswijk et al., 2010). Specifically, we first calculated the average MUA response to all stimulus changes, because it is the optimal estimate of the response in the absence of any rhythm-dependent modulation. We then defined a surrogate input time at 150 ms before the actual input time. We analyzed the LFP phase at the surrogate input time, binned trials according to those phases, and mathematically added the average MUA response across all trials to the average MUA record per phase bin. The resulting additive MUA response component as function of gamma phase is shown as red line in Figure 2C. The additive MUA response component was subtracted from the (total) MUA response to obtain the multiplicative MUA response component (Figure 2D). Phase-dependent modulation depth (MD) of the multiplicative MUA response component was quantified by fitting one cycle of a cosine function (smooth curve in Figure 2D).

The analysis illustrated in Figure 2 for an example MUA recording site was performed for all MUA recording sites of both macaques; in addition, a bias estimate for the cosine fit was obtained for all sites as explained in Experimental Procedures. The observed modulation depths consistently exceeded the bias estimates in the gamma band and also in a band that overlaps with both, the classical alpha and beta bands, and which we therefore address as alpha-beta band (Figure 3). To quantify effect size, we expressed the multiplicative MUA response components as percent of the pre-input MUA rate (Figure 3D). Effect sizes had median values of 20.2% for the alpha-beta band (10-14 Hz) and 21.6% for the gamma band (40-66 Hz) and showed distributions including values exceeding 50% (see Experimental Procedures for quantification of effect sizes).

**Figure 3.**
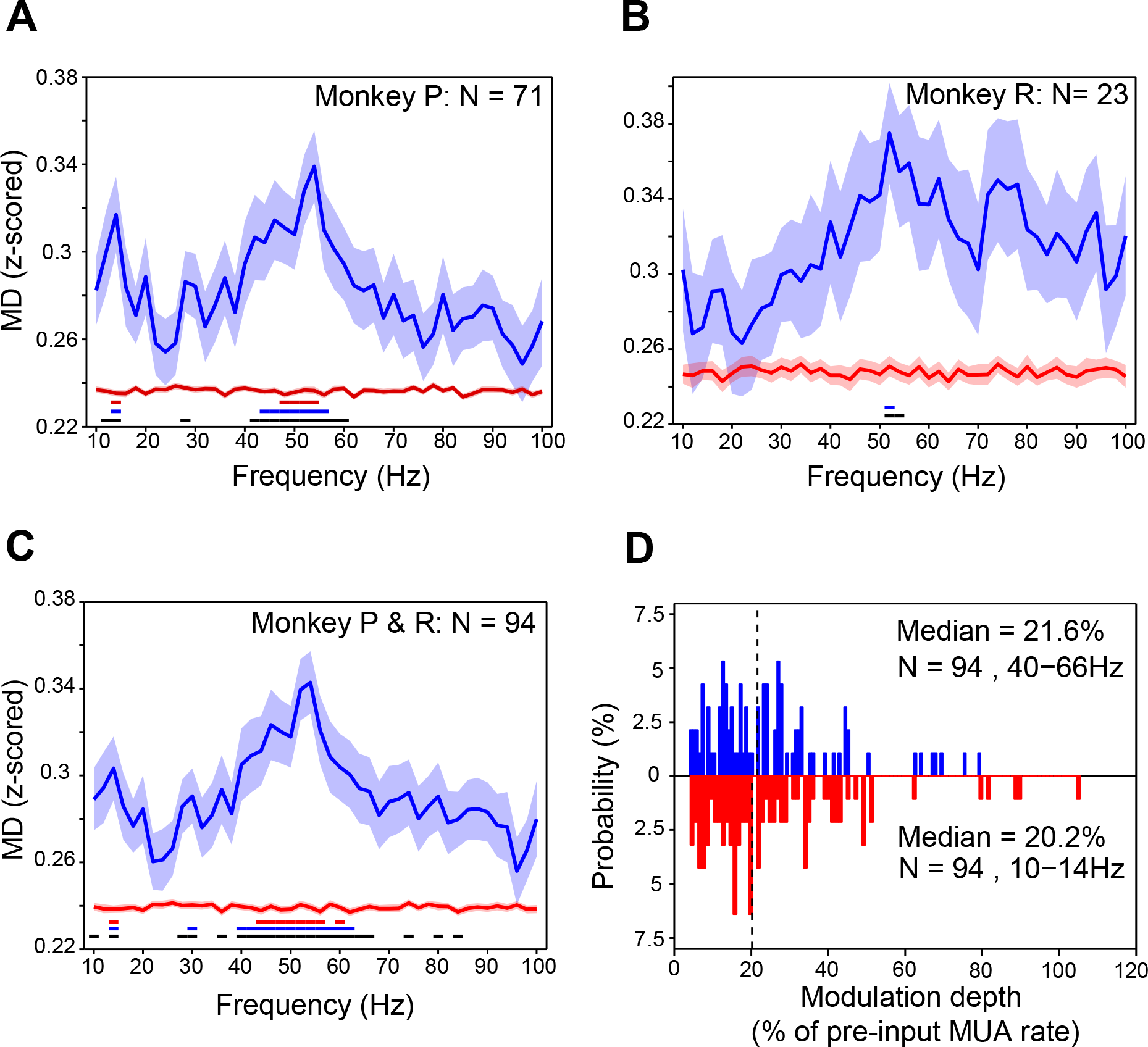
In Macaque V4, Gain Modulation is Prominent for the Gamma Rhythm. (A) Blue curve: Modulation depth of the multiplicative MUA response component as a function of the frequency, for which the pre-input phase was determined. Average over all 71 sites of monkey P after z-transformation per site (see Experimental Procedures). Red curve: Bias estimate. Shaded regions indicate ±1 SEM. Horizontal lines at bottom of plot indicate significance level after correction for multiple comparisons across frequencies: Black lines for p<0.05; blue lines for p<0.01; red lines for p<0.001. (B) Same format as (A), averaged over all 23 sites of monkey R. (C) Same format as (A), averaged over all 94 sites of both monkeys combined. (D) Histogram of modulation depths of the multiplicative MUA response component, expressed as percentage of pre-input MUA rate. Blue histogram on top shows values obtained with binning according to pre-input phase in the gamma-frequency range found significant in (C), i.e. 40-66 Hz; red histogram on bottom shows values obtained with binning according to pre-input phase in the alpha-beta-frequency range found significant in (C), i.e. 10-14 Hz. Dashed vertical lines indicate median values.

### Gamma phase modulates behavioral reaction time

We hypothesized that the phase of visually induced gamma also modulates behavioral reaction times (RTs) in response to the stimulus change. To investigate this, we proceeded similarly to the analysis of MUA responses. Trials were binned according to pre-input LFP phase, and for each phase bin, RTs were averaged. RTs were significantly modulated by the phase of pre-input LFP oscillations between 48 and 52 Hz (Figure 4A, non-parametric permutation test with correction for multiple comparisons across frequencies). On average, LFP phases between 48 and 52 Hz modulated RT by a median of 13 ms (Figure 4B). Thus, the pre-input gamma phase has a small but significant direct influence on behavior.

**Figure 4.**
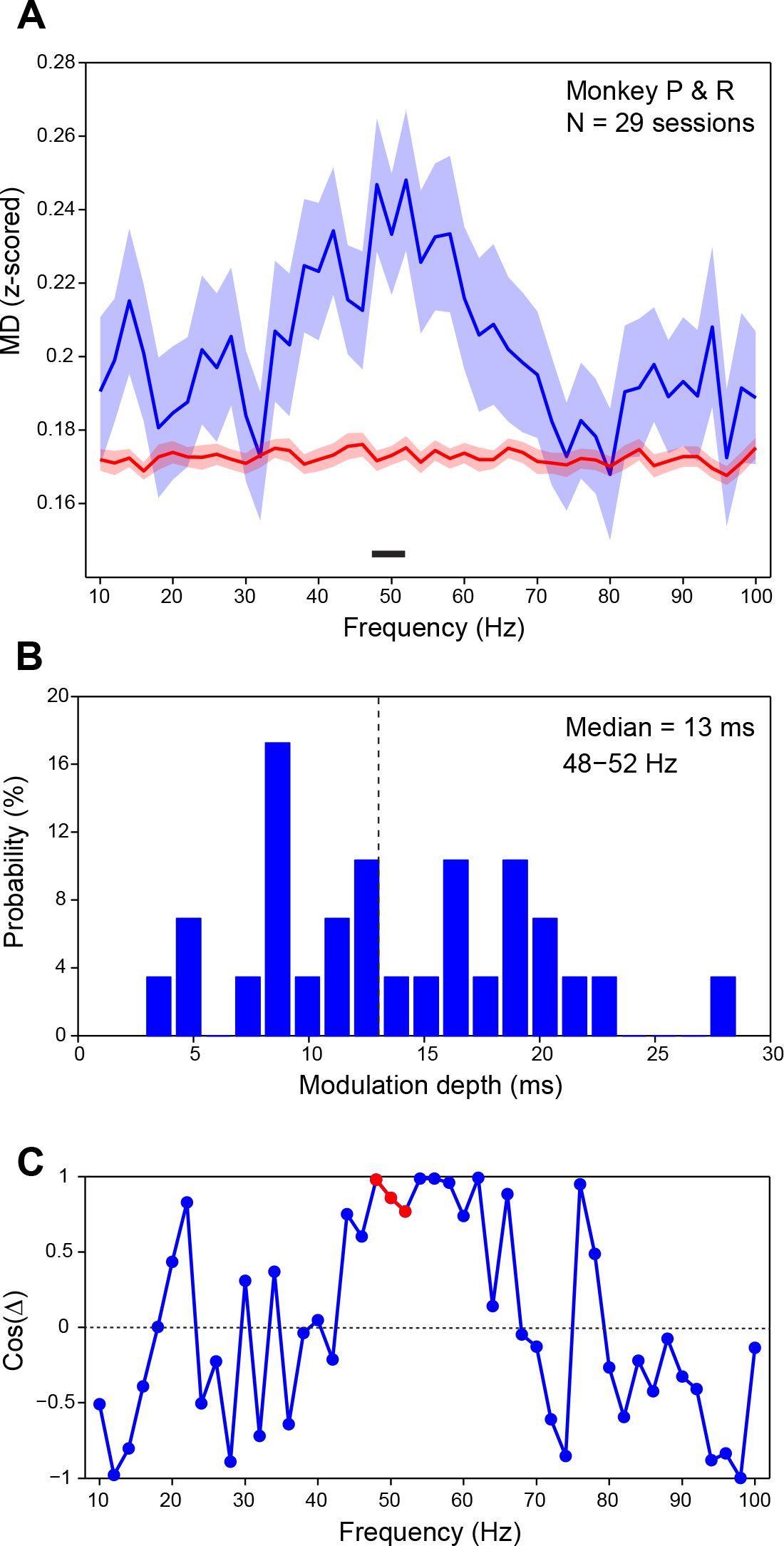
Gamma Phase Modulates Reaction Time, and Similar Gamma Phases Lead to Short Behavioral Reaction Times and Strong Neuronal Responses. (A) Blue: Modulation depth of behavioral reaction times (RTs) by pre-input phase (after z-transformation of RTs per session, by subtraction of mean and division by standard deviation across trials in a session). Red: Bias estimate. Shaded regions indicate ±1 SEM. The black horizontal bar on the bottom indicates significant modulation in the gamma band from 48 to 52 Hz (p<0.05, non-parametric permutation test, corrected for multiple comparisons across frequencies). (B) Distribution of the modulation depths of RTs by pre-input 48-52 Hz phase. (C) The cosine of the difference (Δ) between phases leading to shortest RTs and phases leading to strongest neuronal responses. Cosine values close to one indicate that phases leading to short RTs are close to phases leading to strong neuronal responses. For the frequency range of 48-52 Hz (red dots), phase differences are significantly non-uniform (P=0.03), with an average phase difference of merely 15.3 deg. (A-C) All plots combine the data of both monkeys (N=29 sessions).

So far, we have shown 1.) that pre-input gamma phase partly predicts the MUA response to stimulus change and 2.) that pre-input gamma phase partly predicts behavioral RTs. Therefore, we next investigated whether pre-input gamma phases leading to short RTs are similar to those leading to strong MUA responses. We first selected the MUA recording sites, which showed an individually significant response modulation by pre-input gamma phase (N=69). For those sites, we determined the pre-input gamma phase leading to maximal responses and the pre-input gamma phase leading to shortest RTs (both were determined from the respective cosine function fits). To quantify the similarity between those phases, we took the cosine of the phase difference, which gives a value of 1 for equal phases and a value of −1 for opposite phases. The average cosine spectrum (Figure 4C) shows values close to 1 in the gamma range. In the 48-52 Hz range, for which pre-input phase was significantly predictive of behavioral RT, phase differences were significantly non-uniformly distributed (p=0.03, V-test (Berens, 2009)) with a mean phase difference of merely 15.3 deg. Thus, pre-input gamma phases leading to short RTs also lead to strong MUA responses. This suggests that the influence of gamma phase on MUA responses has functional relevance.

### Effects of the phase of isolated, optogenetically induced gamma

We showed that the phase of ongoing, visually induced gamma in area V4 modulates both, the MUA response to a stimulus change and the corresponding behavioral reaction time. Yet, it remains unclear, whether this effect emerged in V4 or at earlier processing stages. When the visual stimulus induced a gamma rhythm in V4, it most likely induced gamma rhythms also in earlier visual cortical areas, which were at least partly coherent with the V4 gamma. Several previous studies demonstrated visually induced gamma-band coherence between V1 and V4 (Bosman et al., 2012; Brunet et al., 2014; Grothe et al., 2012), and further studies established that gamma in lower visual areas entrains gamma in higher visual areas in a feedforward manner (Bastos et al., 2015a; Bastos et al., 2015b; Bosman et al., 2012; Jia et al., 2013; Michalareas et al., 2016; Roberts et al., 2013; van Kerkoerle et al., 2014). Thus, the effect of V4 gamma phase might emerge at earlier stages. This would be fully in line with our general hypothesis, that a local gamma rhythm modulates the response to randomly timed synaptic test inputs (see Discussion). Nevertheless, we sought to test whether an isolated gamma rhythm could have the same effect. We found that this test could be elegantly performed, when using gamma induced by constant light stimulation of neurons expressing Channelrhodopsin (ChR2). Several previous reports have shown that local ChR2-expressing neuronal populations generate clear gamma rhythms in response to light that is constant or smoothly ramping up (Adesnik and Scanziani, 2010; Akam et al., 2012; Lu et al., 2015). That is, while the light did not contain any temporal structure in the gamma-frequency range, the gamma rhythm was generated by the neuronal network, most likely through reverberant interactions between excitatory and inhibitory neurons (Tiesinga and Sejnowski, 2009; Whittington et al., 2000).

For these experiments, we used the anesthetized cat as a model system (N=2 animals). We injected recombinant adeno-associated viral vectors to express ChR2 in cortical neurons [AAV2.9-CamKIIα-hChR2(H134R)-eYFP]. Vectors were injected into area 21a, the cat homologue of macaque area V4 (Figure 5A) (Payne, 1993). After four to six weeks of expression, recordings were performed under general anesthesia. Subsequently, the animal was perfused and the brain processed histologically. Confocal microscopy showed ChR2-eYFP expression in cortical neurons (Figure 5B). Area 21a recordings showed clear responses to the local application of blue light (473 nm) (Figure 5C). Constant light for a period of 1.25 s induced a pronounced gamma-band rhythm (Figure 5C, middle panel; Figure 6). Simultaneous recordings in area 17, the source of major input to area 21a and the cat homologue of macaque area V1, suggested that optogenetically induced gamma in area 21a did not propagate in the feedback direction to area 17 (Figure 6), consistent with recent reports of the feedforward nature of gamma (Bastos et al., 2015b; Michalareas et al., 2016; van Kerkoerle et al., 2014).

**Figure 5.**
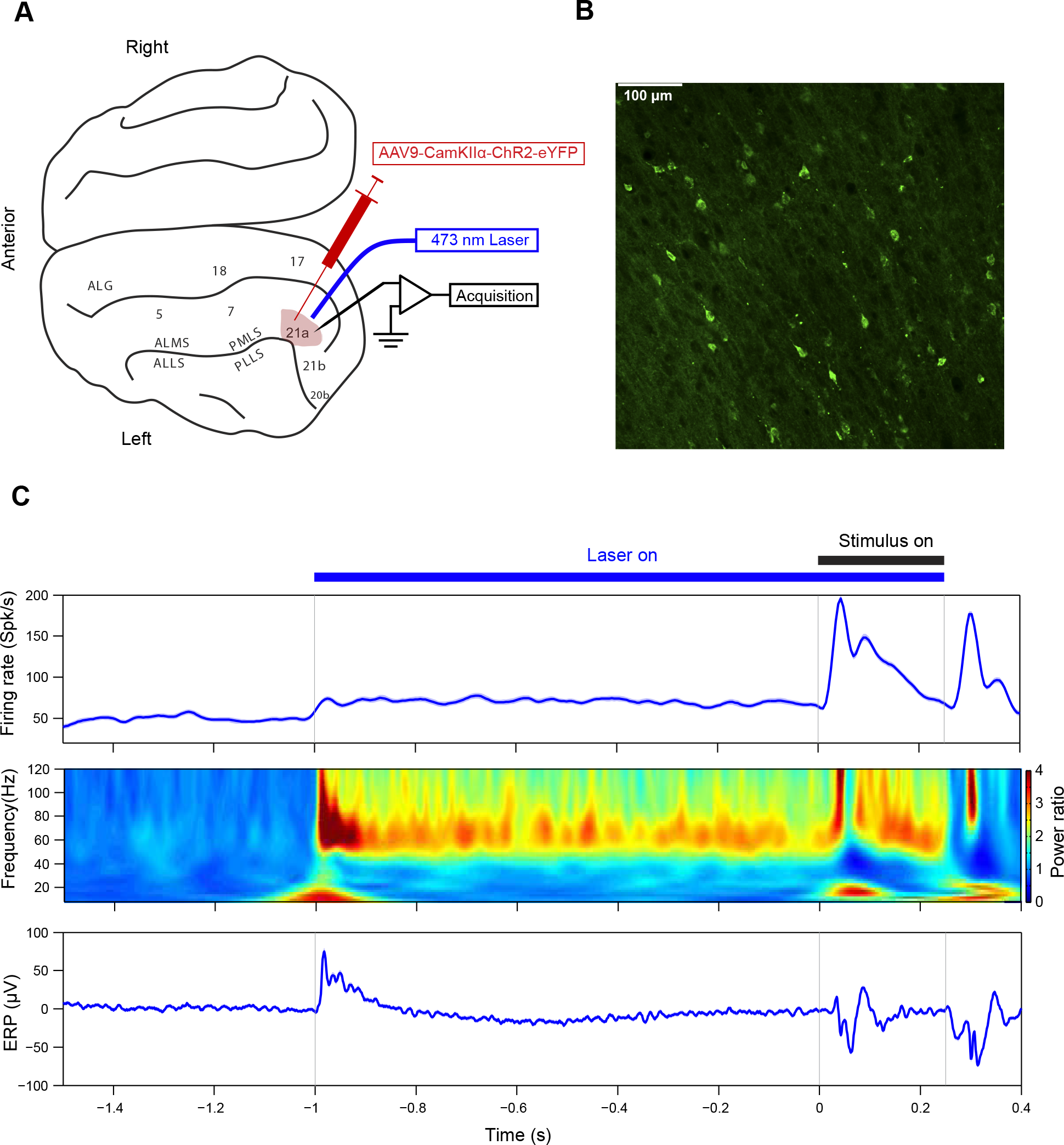
Viral Injection and Expression; Optogenetically and Visually Induced Modulation of MUA Firing Rate, LFP Power and Event-related Potentials in Anesthetized Cat Area 21a. (A) In an initial surgery, the viral vector AAV9-CamKIIα-hChR2(H134R)-eYFP was injected into cat area 21a. After four to six weeks of expression, 473 nm Laser light was applied through a fiber placed above area 21a, visual stimuli were shown and electrophysiological recordings performed from area 21a. (B) Example histological section, showing the distribution of eYFP-labeled neurons in area 21a through fluorescence microscopy. (C) Responses of one example recording site during optogenetic and visual stimulation as indicated by the horizontal lines above the top panel. Laser stimulation commenced first, followed one second later by visual stimulation. Top panel: MUA firing rate. Middle panel: LFP power. Bottom panel: Event-related potential. The shaded regions around the lines in the top and bottom panels indicate ±1 SEM; they are hardly visible behind the actual lines.

**Figure 6.**
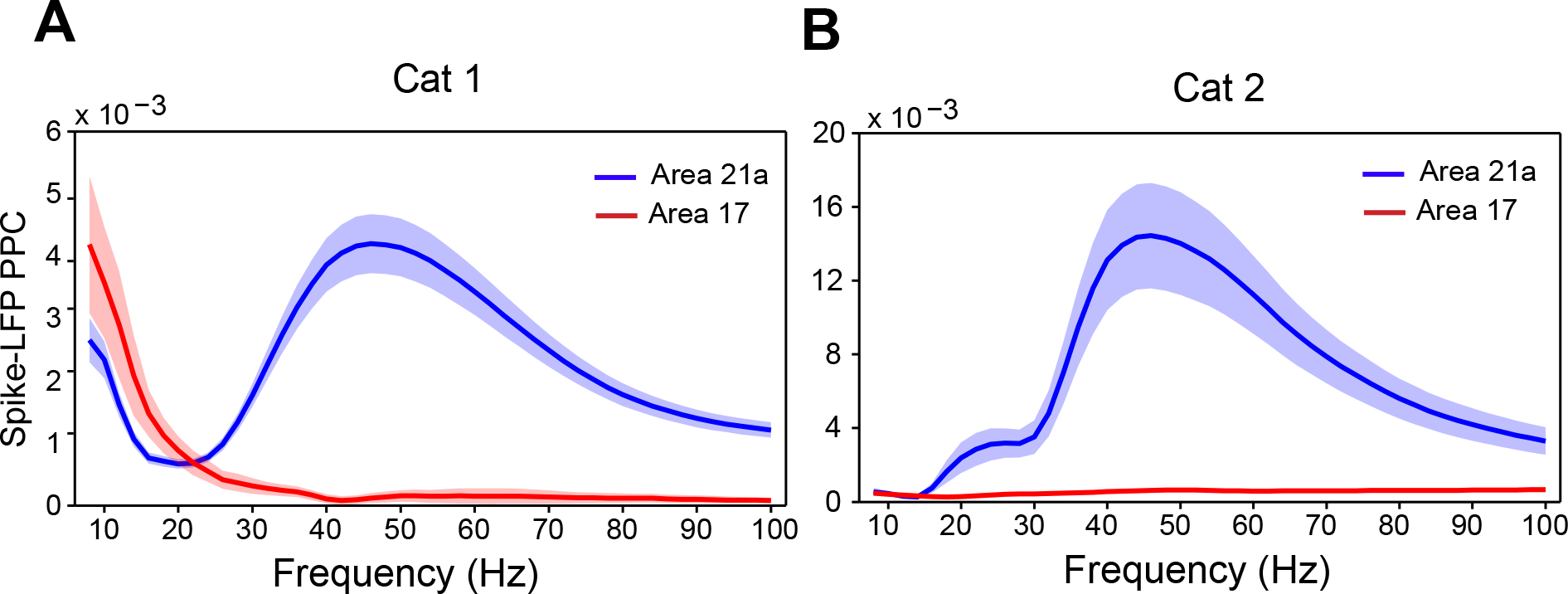
Optogenetic Stimulation of Area 21a Induces Gamma in Area 21a and Not in Area 17. (A)Spike-LFP locking in area 21a (blue) and area 17 (red) during optogenetic stimulation of area 21a in the absence of visual stimulation. Each line shows the average over all respective recording sites of cat 1 (Area 21a: N=57; area 17: N=11). Shaded regions indicate α1 SEM. (B) Same as (A), but showing the averages over all respective recording sites of cat 2 (Area 21a: N=33; area 17: N=38).

Having established this optogenetically induced gamma in area 21a, we produced synaptic test inputs through visual stimulation. Visual stimuli were presented at 1 s after onset of optogenetic stimulation and lasted for 0.25 s. We analyzed the MUA response to visual stimulus onset as a function of the LFP phase prior to the input time. We used the same approach as in the analysis of the macaque V4 data, with one difference: Whereas the visually induced gamma peak frequencies in the two macaques happened to be almost identical, the optogenetically induced gamma peak frequencies differed across cats and recording sessions (Figure S1). These differences were probably due to differences in local density of opsin expression, effective light intensity, the state of the anesthetized cat or combinations of those factors. Therefore, we determined the optogenetically induced gamma peak frequency per recording site and aligned the analysis to it. The mean input times were 27.7±0.3 ms (Cat 1) and 32.3±0.9 ms (Cat 2). We found that gamma induced locally in area 21a by optogenetic stimulation was sufficient to multiplicatively modulate the MUA response to visual stimulus onset (Figure 7A–C). The median effect size was 21.6% (Figure 7D).

**Figure 7.**
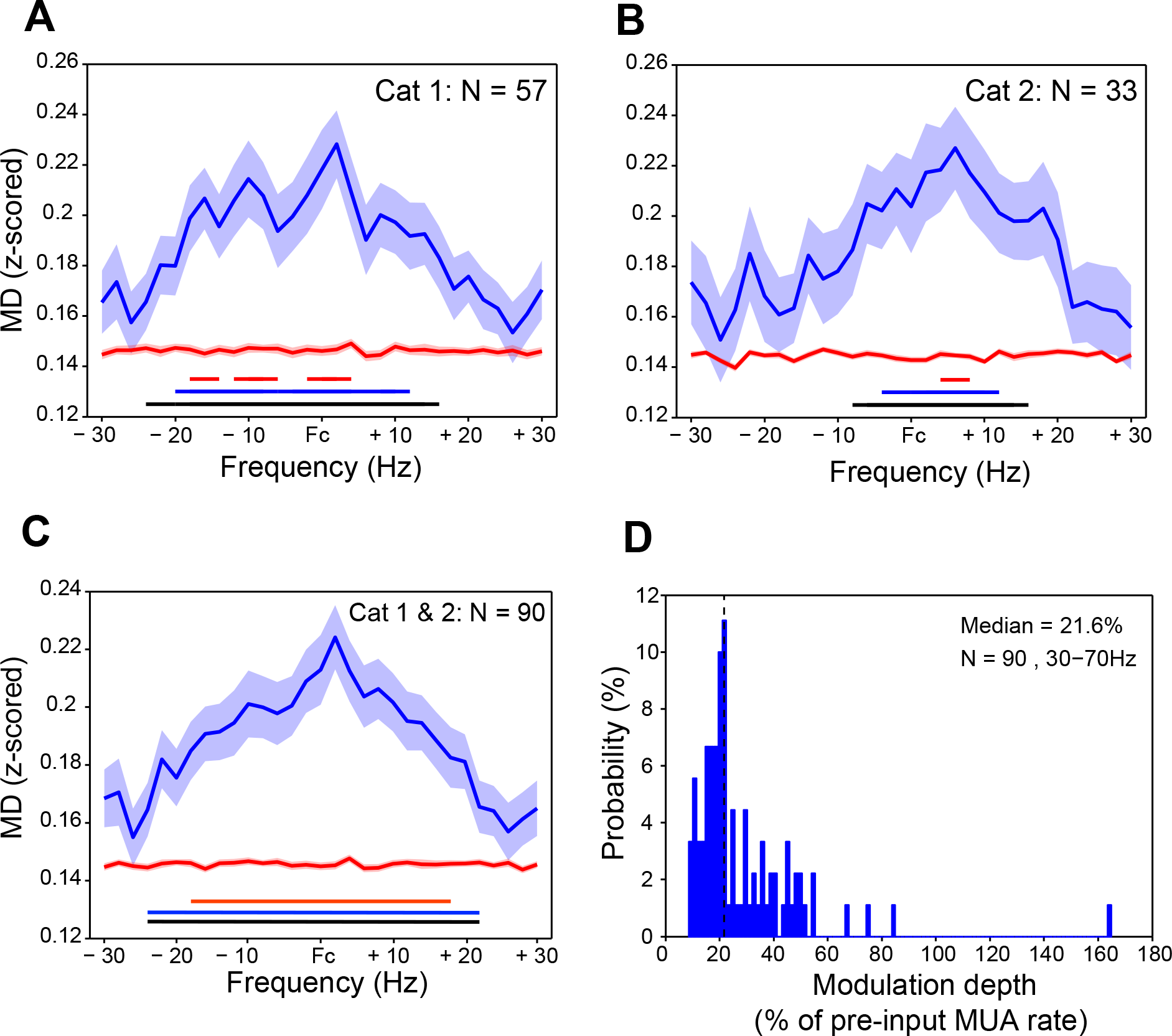
Gain Modulation by Optogenetically Induced Gamma. (A) Blue curve: Modulation depth of the multiplicative MUA response component as a function of the frequency, for which the pre-input phase was determined. Average over all 57 recording sites in area 21a of cat 1 after z-transformation per site (see Experimental Procedures). Per recording site, the spectral analysis was aligned to the gamma peak frequency induced at that site by optogenetic stimulation (see Figure S1). Red curve: Bias estimate. Shaded regions indicate ±1 SEM. Horizontal lines at bottom of plot indicate significance level after correction for multiple comparisons across frequencies: Black lines for p<0.05; blue lines for p<0.01; red lines for p<0.001. (B) Same format as (A), averaged over all 33 area 21a sites of cat 2. (C) Same format as (A), averaged over all 90 area 21a sites of both cats combined. (D)Histogram of modulation depths of the multiplicative MUA response component, expressed as percentage of pre-input MUA rate. Dashed vertical line shows median.

## Discussion

We found that gamma-band activity rhythmically modulates multiplicative response gain. Gain modulation exceeded 50% in some cases and had a median size of ~20%, which is likely an underestimation as explained below. The findings held across gamma induced in awake macaque V4 by sustained visual stimulation and gamma induced in anesthetized cat area 21a by constant optogenetic stimulation. The awake macaque data allowed us to investigate the relevance of gamma phase for behavior. This showed that gamma phases leading to short behavioral reaction times were similar to gamma phases leading to strong MUA responses. Optogenetic stimulation allowed us to investigate the effect of “isolated” gamma in a higher visual area, that was most likely not preceded by substantial gamma in lower visual areas. This showed that such isolated optogenetically induced gamma leads to multiplicative gain modulation of similar size as visually induced gamma.

In a previous paper, we provided first evidence that effective connectivity between two visually driven local neuronal groups in awake cat or monkey visual cortex depends on the phase relation between the respective local gamma rhythms (Womelsdorf et al., 2007). For each pair of recording sites, we segmented the data into 0.25 s long epochs, estimated the phases of local neuronal rhythms and sorted epochs into six bins, according to their phase relations. Per frequency, for which the phase relation was estimated, we quantified effective connectivity as the correlation between the respective power values, across the epochs of a given phase-relation bin. We found that effective connectivity depended systematically on the phase relation, primarily in the gamma-frequency band. A subsequent mathematical modeling study simulated two gamma-synchronized neuronal groups and found that the gamma phase relation between the groups does not only determine their power correlation, but also their mutual transfer entropy (Buehlmann and Deco, 2010). Transfer entropy is an information theoretical measure that quantifies the statistical dependence between systems and is able to distinguish between driving and responding elements and therefore between shared and transmitted information. A recent experimental study investigated transfer entropy between multiple simultaneous recordings in anesthetized macaque area V1 (Besserve et al., 2015). Transfer entropy was influenced by the phase relation between local gamma-band rhythms. In particular, dynamic changes in the stimulus led to directed gamma-band waves and a relative increase in the amount of information flowing along the instantaneous direction of the gamma wave.

While these studies together strongly suggest that the phase relation among gamma-band rhythms affects the strength and direction of influences between the respective neuronal groups, the possibility remains that those phase relations are not the cause but the consequence of the neuronal influences. It is conceivable that other mechanisms modulate effective connectivity, and that enhanced effective connectivity subsequently leads to particular phase relations. The current results provide compelling evidence that the gamma rhythm is actually a cause of modulations in effective connectivity. We show that the gain is modulated within each gamma cycle, as a function of gamma phase. If this gamma-rhythmic gain modulation were due to a mechanism other than the gamma rhythm itself, this mechanism would necessarily oscillate at the relatively high gamma frequency and in synchrony with the gamma rhythm (without actually being the gamma rhythm). While such arbitrarily complex assumptions can explain essentially any set of results, it is much more parsimonious and physiologically plausible that the gamma rhythm itself modulates gain rhythmically and thereby enhances the effective connectivity of inputs synchronized to gamma phases of high gain.

Our central experimental approach has been to assess the response to a temporally unpredictable visual event, i.e. a stimulus change in the monkey recordings and a stimulus onset in the cat recordings. Because the stimulus event is physically identical in all trials, the strength of the resulting synaptic input should be constant, at least at the earliest stages of visual processing (with the exception of uncontrolled fluctuations arising from physiological noise). Our finding that physically identical stimulus events lead to varying postsynaptic responses that depend systematically on the pre-input gamma phase, unequivocally demonstrates the rhythmic modulation of postsynaptic gain. Importantly, we cannot conclude from our measurements that the postsynaptic gain modulation emerged in the very neurons, from which we recorded the spiking activity. In the awake macaque experiments, the postsynaptic gain modulation might have emerged in any neuron on the way from the retina to the recorded V4 neurons. Yet, wherever the modulation emerged, it there constituted a postsynaptic modulation, because of our use of identical stimulus events across trials. Note that any modulation that emerged at an earlier stage would become visible in our analysis only, if the gamma phase at this earlier stage were coherent with the gamma phase recorded in V4. Gamma coherence between early and intermediate level visual areas is clearly present but of small magnitude (Bastos et al., 2015a; Bosman et al., 2012; Grothe et al., 2012) and typically not transitive across multiple processing stages (Zandvakili and Kohn, 2015). Thus, local gamma rhythms in V4 and several preceding visual areas likely lead to local gain modulation, and the gain modulation that we observed as a function of V4 gamma phase likely emerges fully or largely in V4. This interpretation is consistent with the results of the optogenetics experiments. Optogenetic stimulation induced a gamma rhythm in area 21a and no appreciable gamma rhythm in area 17. The modulation of gain by the area 21a gamma phase was similar to the modulation found in the awake macaque. We note that it is not possible to fully equate strength and extent of optogenetically and visually induced gamma. Nevertheless, visually and optogenetically induced gamma were of similar magnitude and so were the corresponding modulations of gain.

While visually and optogenetically induced gamma rhythms led to gain modulation of similar size, our size estimates likely remain below the actual physiologically relevant gain modulation. We consider the physiological relevance of gamma-phase dependent gain modulation primarily in the context of the Communication-through-Coherence (CTC) hypothesis. The CTC hypothesis posits that a local gamma rhythm modulates the gain of individual synaptic inputs. Therefore, the ideal test would have been to deliver individual synaptic inputs to a neuron embedded in a gamma-rhythmic network and to quantify the degree to which the gamma phase modulates the probability with which the synaptic input is followed by a postsynaptic spike. We deem it likely that gain modulation for individual synaptic inputs is larger than the gain modulation found here for large barrages of synaptic inputs triggered by visual stimulus events. Synaptic input barrages likely override part of the gain-modulating effect of the gamma rhythm and create a partial ceiling effect. Future experiments will need to aim at delivering smaller barrages or in fact individual synaptic inputs to neurons embedded into in-vivo gamma-rhythmic networks, similar to previous in-vitro experiments (Volgushev et al., 1998). This is challenging, because the strength of the test inputs must not change as a function of gamma phase, because this would constitute a presynaptic modulation, confounding the hypothesized postsynaptic modulation. Another reason we likely underestimated the effect size is noise in the recording and estimation of the relevant gamma phase. The gamma phase, that is actually relevant in the CTC context, stems from the gamma rhythm that impacts postsynaptic spiking, i.e. the gamma rhythm in the membrane potential fluctuations of the neurons, from which we recorded the spike response. In vivo and particularly in the awake macaque, these fluctuations can best be approximated by the LFP (Haider et al., 2016). Yet, the LFP contains substantial measurement noise and additional physiological components, reflecting e.g. current flows in more distant, functionally separate neurons, resulting in gamma-phase noise. In addition, there is substantial noise in the form of uncertainty in the estimation of the gamma phase at the time when it putatively interacts with synaptic input, i.e. the “input time”. The gamma phase at the input time can only be estimated by analyzing a finite-length window preceding the input time. The finite-length window is necessary to obtain specificity for a frequency band. Thus, our spectrally defined phase reflects a time window rather than a time point, and, because the gamma rhythm has a short autocorrelation length, this only approximates the phase at the input time point. A particular challenge in the present context was that estimating the phase at the input time had to exclude data recorded after the input time. Spectral phase estimation requires, to avoid aliasing, the application of a data taper that approaches zero at its edges, which would reduce the weight of data from close to the input time. To avoid this, we extrapolated the data by means of an autoregressive model before spectral estimation. Yet, even if this approach was the best possible given the constraints, it could not avoid substantial estimation noise that is expected to reduce the observed effect size. These considerations assume that the synaptic test inputs arrived and interacted with the ongoing oscillation at one time point, whereas they likely arrived with a certain temporal distribution. This distribution must have been short relative to the gamma cycle, because otherwise any gain modulation effect would have been averaged out across the different gamma phases, over which input had been spread out. Yet, some temporal spread of the synaptic inputs driven by the stimulus events is very likely, and this spread further led to underestimation of the true effect size.

The likely temporal spread of stimulus-event-related synaptic inputs also precluded a simple interpretation of the absolute gamma phase leading to maximal peak responses. We determined this phase relative to the input time, which we conservatively defined as the last time bin before LFP inter-trial coherence (ITC) deviated significantly from pre-stimulus-event values. While this approach safely excludes post-input data from the estimation of the pre-input phase, it introduces a certain delay between the estimated input time and the actual temporal distribution of synaptic test inputs. This delay likely differed slightly across the different MUA clusters, e.g. due to uncertainty in ITC onset estimation and to actual physiological differences in the temporal spread of synaptic inputs. Even a delay of merely 5 ms will result in a phase rotation at 50 Hz of 90 degree. Therefore, in order to interpret the phases leading to maximal peak responses, we compared them to the phases leading to shortest behavioral reaction times. Despite the substantial noise and uncertainty involved on both sides, this analysis revealed that, across different MUA recordings sites, gamma phases leading to shortest reaction times were close to gamma phases leading to strongest MUA responses. This suggests that the observed gamma phases leading to maximal responses are actually meaningful and that the gain modulation by the gamma rhythm has direct behavioral relevance.

## Experimental Procedures

Experiments were performed on two awake macaque monkeys and on two anesthetized cats. Data analysis for the two datasets followed the same approach.

### Experiments on macaques

Experiments were performed on two adult macaque monkeys, following the guidelines of the National Institutes of Health. Recordings were performed in area V4, while animals were awake and performing a selective visual attention task. The data analyzed here have been used in previous studies (Bosman et al., 2009; Brunet et al., 2014; Buffalo et al., 2011; Buffalo et al., 2010; Fries et al., 2001; Fries et al., 2008; Liang et al., 2005; Maris et al., 2013; Vinck et al., 2013; Womelsdorf et al., 2006; Womelsdorf et al., 2007).

### Visual stimulation and behavioral task

Visual stimulation, receptive field mapping and attentional task are described in detail in (Fries et al., 2008), and we report here only the essential points. Stimuli were presented on a 17 inch cathode ray tube monitor 0.57 m from the monkey’s eyes with a refresh rate of 120 Hz non-interlaced. Stimulus generation and behavioral control were accomplished with the CORTEX software package (http://dally.nimh.nih.gov). The orientation of the drifting grating placed inside the receptive fields was selected so that it maximally co-activated the simultaneously recorded units. A second grating patch, rotated by 90 deg and otherwise identical, was placed outside the receptive fields.

Several slightly different trial structures were used with different attentional cueing regimes (trial-by-trial cueing using as cue either short lines or the fixation point color, or trial-block cueing). As the attentional cueing regime is not relevant for the present analysis, we describe here the general trial structure. A trial started when the monkey touched a bar and directed its gaze within 0.7° of the fixation spot. After a baseline period of at least 1.5 s, the stimuli were presented, one cued as target, the other as distracter. Either the target or the distracter (equal probability) changed color (from black/white to black/yellow) at an unpredictable moment between 0.5 and 5 s after stimulus onset (flat random distribution of change times across trials). If the distracter changed first, the target changed later, between the distracter change time and 5 s post stimulus onset. If the monkey released the bar within 0.15-0.65 s of a target change, a fluid reward was given. If the monkey released within the same time period after a distracter, a timeout was given. Trials were aborted if the monkey broke fixation or released the bar prematurely. In a typical recording session, monkeys completed 200 to 600 correctly performed trials.

### Neurophysiological recordings

Magnetic resonance imaging was used to localize the prelunate gyrus. Recording chambers were implanted over the prelunate gyrus under surgical anesthesia. In each recording session, three to four tungsten microelectrodes (impedances around 1 MΩ at 1kHz) were advanced separately through the intact dura at a very slow rate (1.5 μm/s) to minimize deformation of the cortical surface by the electrode (“dimpling”). Electrodes were horizontally separated by 650 or 900 μm. Standard electrophysiological techniques (Plexon MAP system) were used to obtain MUA and LFP recordings. For MUA recordings, the signals were filtered with a passband of 100 to 8000 Hz, and a threshold was set interactively to retain the spike times of small clusters of units. For LFP recordings, the signals were filtered with a passband of 0.7 to 170 Hz and digitized at 1 kHz.

### Experiments on cats

Two adult female domestic cats were used. All procedures complied with the German law for the protection of animals and were approved by the regional authority (Regierungspräsidium Darmstadt). After an initial surgery for the injection of viral vectors and a 4-6 week period for virus expression, recordings were obtained during a terminal experiment under anesthesia.

### Viral vector injection

For the injection surgery, anesthesia was induced by intramuscular injection of Ketamine (10 mg/kg) and Medetomidine (0.02 mg/kg), cats were intubated, and anesthesia was maintained with N_2_O:O_2_ (60/40%), isoflurane (~1.5%) and Remifentanil (0.3 μg/kg/min). A rectangular craniotomy was made over the left hemisphere (AP: 0 to −8 mm, ML: 9 to 15 mm), area 21a was identified by the pattern of sulci and gyri, and the dura was removed over part of area 21a. Four injection sites in area 21a were chosen, avoiding major blood vessels, with horizontal distances between injection sites of at least 1 mm. At each site, a Hamilton syringe (34G needle size; World Precision Instruments) was inserted under visual inspection to a cortical depth of 1 mm below the pia mater. Subsequently, 2 μl of viral vector solution (AAV2.9-CamKIIα-hChR2(H134R)-eYFP; titer 1.06e13 GC/ml; Penn Vector Core, Philadelphia, PA) was injected at a rate of 150 nl/min. After each injection, the needle was left in place for 10 min before withdrawal to avoid reflux. Upon completion of injections, the dura opening was covered with silicone foil and a thin layer of silicone gel, the trepanation was filled with dental acrylic, and the scalp was sutured.

### Neurophysiological recordings

For the recording experiment, anesthesia was induced and initially maintained as during the injection surgery, only replacing intubation with tracheotomy and Remifentanyl with Sufentanil. After surgery, during recordings, isoflurane concentration was lowered to 0.6%−1.0%, eye lid closure reflex was tested to verify narcosis, and Vecuronium (0.25 mg/kg/h i.v.) was added for paralysis during recordings. Throughout surgery and recordings, Ringer solution plus 10% Glucose was given (20 ml/h during surgery; 7 ml/h during recordings), and vital parameters were monitored (ECG, body temperature, expiratory gases).

Each recording experiment consisted of multiple sessions. For each session, we inserted either single or multiple tungsten microelectrodes (~1 MΩ at 1 kHz, FHC), or three to four 32-contact probes (100 μm inter-site spacing, ~1 MΩ at 1 kHz; NeuroNexus or ATLAS Neuroengineering) in area 21a. In some sessions, an additional 3-4 of the same 32-channel probes were inserted into area 17. Standard electrophysiological techniques (Tucker Davis Technologies system) were used to obtain MUA and LFP recordings. For MUA recordings, the signals were filtered with a passband of 700 to 7000 Hz, and a threshold was set interactively to retain the spike times of small clusters of units. For LFP recordings, the signals were filtered with a passband of 0.7 to 250 Hz and digitized at 1017 Hz.

Optogenetic stimulation was done with a 473 nm (blue) laser or with a 470 nm (blue) LED (Omicron Laserage, Germany). A 597 nm (yellow) laser was used as control in some sessions and did not induce gamma-band activity. Laser light was delivered to cortex through a 200 μm diameter multimode fiber, LED light through a 2 mm diameter multimode fiber. Fiber endings were placed just next to the recording sites with a slight angle relative to the electrodes. Illumination was applied to the recorded patch of area 21a for 1.25 s at a constant level. Intensity was titrated to induce clear gamma-band activity and totaled 1-10 mW when measured at the fiber ending. At 1 s after illumination onset, a visual stimulus was presented on a liquid crystal display (LCD, Samsung 2233RZ) with a screen update frequency of 120 Hz. Contact lenses were placed into the two eyes to equate their refraction as well as possible. The eye-to-screen distance was determined by the mean refraction index of the two eyes with their respective contact lenses. If necessary, prisms were used to align the eyes. Visual stimuli were presented for 0.25 s. They were either a static bar or a static grating patch inside the RFs of the recorded neurons, or a static full-field grating. Stimulus generation and control used Psychtoolbox-3, a toolbox in MATLAB (MathWorks, Natick, MA)(Brainard, 1997).

### Histology

After the experiment, the animal was euthanized with pentobarbital sodium and transcardially perfused with phosphate buffered saline (PBS) followed by 4% paraformaldehyde. The brain was removed, post-fixed in 4% paraformaldehyde and subsequently soaked in 10%, 20% and 30% sucrose-PBS solution, respectively until the tissue sank. The cortex was sectioned in 50 μm thick slices. The slices were investigated with a confocal laser microscope (Nikon Instruments) for eYFP-labelled neurons.

## Data analysis

### Spike densities, power spectra, ERPs and spike-LFP PPCs

MUA was smoothed with a Gaussian kernel (SD = 12.5 ms, truncated at ±2 SD) to obtain the spike density.

The LFP power spectra shown in Figures 1 and 5 were calculated with windows that were adjusted for each frequency to have a length of 4 cycles. Those windows were moved across the data in steps of 1 ms. For each frequency and window position, the data were Hann tapered, Fourier transformed, squared and divided by the window length to obtain power density per frequency. These power values were then expressed as percent change of the average power in the baseline, −0.5 to −0.25 s before onset of the visual stimulus in the macaque recordings, and before onset of optogenetic stimulation in the cat recordings. Finally, power-change values were averaged over all recording sites.

ERPs were calculated as time-domain LFP averages after baseline subtraction.

Spike-LFP locking was quantified by calculating the spike-LFP PPC (pairwise phase consistency) (Vinck et al., 2010), a metric that is not biased by trial number, spike count or spike rate. Spike and LFP were always taken from different electrodes to avoid contamination of the LFP by low-frequency components of the spike wave shape. For PPC calculation, the LFP around each spike in a window of 4 cycle length was Hann tapered and Fourier transformed. Spike train history effects were avoided by using exclusively pairs of spikes from separate trials. For a given MUA channel, spike-LFP PPC was calculated relative to all available LFPs and then averaged.

### Input time and pre-input LFP phase

We investigated whether the LFP phase just before the time of synaptic test inputs (driven by stimulus change in the macaque and by stimulus onset in the cat) predicts the later MUA response. To estimate the time of synaptic inputs, the “input time”, we used the LFP, because it provides a sensitive metric of the bulk synaptic inputs to the local neuronal group. We reasoned that the first significant stimulus-related response in the LFP should occur shortly after synaptic input arrives and it might in fact reflect the synaptic input directly. As a particularly sensitive metric of LFP response onset, we calculated the inter-trial coherence (ITC).

We needed to estimate the LFP phase as close as possible to the input time, while excluding any influence from after the input time. We created a bank of second-order band-pass Butterworth filters, with passband frequencies spaced between 10 and 100 Hz in steps of 2 Hz. Passband width scaled with passband frequencies, such that the lower (upper) cutoff was always at the passband frequency (F) minus (plus) F/8 Hz. The LFP starting from 0.4 s before the input time was filtered with this filter bank in the forward direction only. Subsequently, LFPs were downsampled to 250 Hz. Per LFP signal and per filter frequency, an autoregressive (AR) model of order 6 was fitted separately to each trial and then averaged over trials. The AR model was used to extrapolate the signal 4 cycles beyond the input time, which was done to avoid edge artifacts of the subsequent Hilbert transform (Chen et al., 2013). The Hilbert transform provided the analytic signal, from which the phase at the last (1 ms) sample before the input time was obtained, which we defined as the pre-input phase.

### MUA responses to stimulus-driven input and their modulation by pre-input phase

We investigated the effect of the pre-input phase on the MUA response to stimulus events, i.e. stimulus changes in macaque V4 and stimulus onsets in cat area 21a. Per recording site, peri-event MUA spike densities were averaged over all trials, and a Gaussian function was fitted, whose mean was used as MUA peak response time for that recording site. Per trial, spike densities from 5 ms before to 5 ms after the MUA peak response time were averaged to obtain the MUA response.

Per recording site and per frequency, trials were grouped according to the phase at the input time into 6 phase bins centered at plus and minus 30, 90 and 150 degrees, respectively (see Figure 2B for illustration). For each phase bin, a number of trials with phases closest to the phase-bin center were chosen. In macaques, this number was 75 trials, in cats it was 200 trials. MUA responses were averaged over the trials in a given phase bin.

A dependence of the MUA response on pre-input gamma phase might be due to a simple additive superposition of a constant MUA response onto the ongoing gamma-modulated MUA firing. Therefore, as explained in the main text, we obtained, per recording site, frequency and phase bin, an estimated additive MUA response component. We subtracted the additive MUA response component from the (total) MUA response (of that recording site, frequency and phase bin) to quantify the respective multiplicative MUA response component. To combine multiplicative MUA response components across recording sites, a z-transformation was done per recording site, by subtracting the mean and dividing by the standard deviation of the total MUA response across trials.

The phase-dependent modulation of the z-transformed multiplicative MUA response components was quantified by fitting one cycle of a cosine function and defining the peak-to-peak amplitude as modulation depth (MD) (Figure 2D). We fitted both the cosine amplitude and phase to avoid strong assumptions about the phase leading to the strongest response. Because cosine fits without pre-determined phase always result in positive modulation depth, we estimated this bias. We randomly combined phases with z-transformed multiplicative MUA components and repeated the cosine fit 100 times. The average MD across those 100 randomizations is the bias estimate and is shown in Figures 3 and 7 as red line.

To quantify the size of the phase-dependent modulation of the (non z-transformed) multiplicative MUA response component, i.e. to quantify effect size (Figure 3D, 7D), we used the modulation depths without subtraction of the bias. The bias is due to the fact that even noisy, i.e. random, variations in the multiplicative MUA response component will lead to a non-zero amplitude of the fitted cosine function. Importantly, those noisy variations are expected to randomly increase or decrease the multiplicative MUA response component, i.e. they are not expected to add to the true multiplicative MUA response component in a way that would systematically change the observed multiplicative MUA response component. This might appear counterintuitive given that we had to statistically test the observed multiplicative MUA response component against the bias estimate. To illustrate the situation, we would like to draw the analogy to extracellular spike recordings in the presence of the typical high-frequency noise. Spikes of a given neuron become visible, when their amplitude exceeds the noise level. Yet, quantification of the peak-to-peak amplitude of the average spike waveform does not subtract the noise, because the noise superimposes with the spike randomly in a positive and negative manner and does not systematically change spike amplitude.

### Statistical testing

Per recording site and per frequency, we obtained the observed modulation depth (MD) and the corresponding bias estimate, i.e. per site, we obtained an MD spectrum and a bias spectrum. We tested whether those spectra differed consistently across recording sites. We calculated paired t-tests between MD and bias spectra, across sites. Statistical inference was not based directly on the t-tests (and therefore corresponding assumptions will not limit our inference), but merely the resulting t-values were used as difference metric for the subsequent non-parametric permutation test. For each of 10000 permutations, we did the following: We made a random decision per site to either exchange the MD spectrum and the bias spectrum or not; we performed the t-test; we placed the largest t-value across all frequencies into the randomization distribution; this latter step implements multiple comparison correction across frequencies (Nichols and Holmes, 2002). Finally, we compared the observed t-values with the randomization distributions to derive p-values for a twosided test, corrected for the multiple comparisons across frequencies. All analyses were done with MATLAB and the Fieldtrip toolbox (Oostenveld et al., 2011).

## Author contributions

Conceptualization, J.N. and P.F.; Methodology, J.N., T.W., C.M.L., I.D. and P.F.; Software, J.N., T.W., and C.M.L.; Formal Analysis, J.N.; Investigation, J.N., T.W., C.M.L., and P.F.; Writing-Original Draft, J.N. and P.F.; Writing – Review & Editing, J.N., T.W., C.M.L., R.D., I.D. and P.F.; Supervision, I.D., R.D., and P.F.; Funding Acquisition, I.D., R.D., and P.F.

## Acknowledgments

The authors thank Jarrod Dowdall, Dmitriy Lisitsyn and Craig Richter for helpful suggestions on phase estimation using AR models, Alina Peter, Georgios Spyropoulos and Martin Vinck for helpful suggestions on data analysis, and Iris Grothe and Wolf Singer for helpful suggestions on the text. PF acknowledges grant support by DFG (SPP 1665, FOR 1847, FR2557/5-1-CORNET), EU (HEALTH-F2-2008-200728-BrainSynch, FP7-604102-HBP, FP7-600730-Magnetrodes), a European Young Investigator Award, NIH (1 U54MH091657-WU-Minn-Consortium-HCP), and LOEWE (NeFF). ID acknowledges grant support by DFG (SPP 1665), EU (ERC Starting Grant OptoMotorPath), BMBF (Bernstein Award 2012), LOEWE (NeFF), and the Minna-James-Heinemann Foundation.

## Supplemental Information

**Figure S1.**
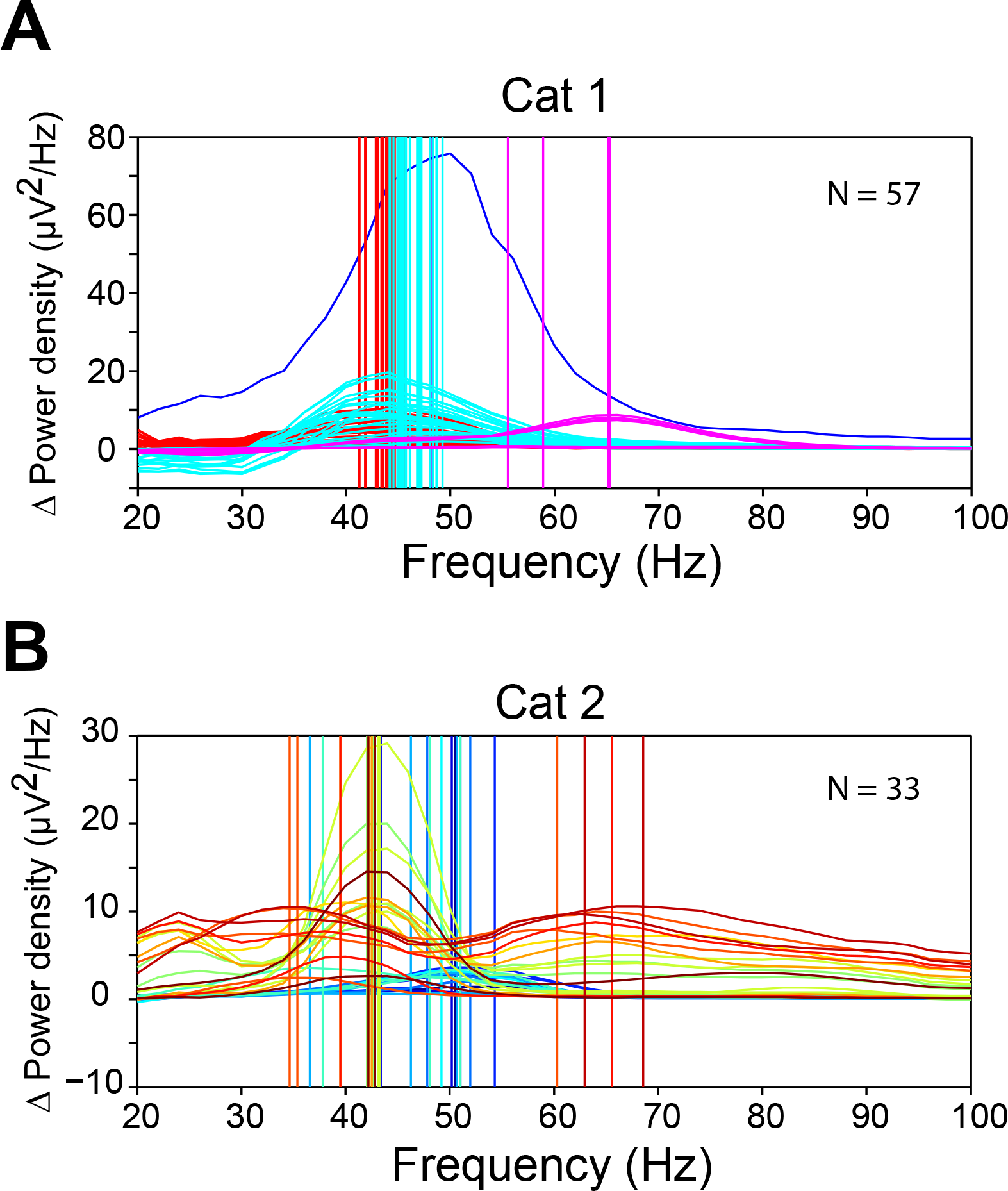
Related to Figure 7. Optogenetically Induced Gamma Shows Variable Peak Frequencies. (A) Difference (Δ) between power during optogenetic stimulation and pre-stimulation baseline. Each spectrum shows data from one of the 57 recording sites in area 21a of cat 1. For each recording site, a vertical line is drawn at the gamma peak frequency, which was determined by a Gaussian fit. Spectra and vertical lines for simultaneously recorded sites are shown in the same color. (B)Same format as (A), but each line shows data from one of the 33 recording sites in area 21a of cat 2.

